# AI-Informed neoantigen prioritization enables a multi-epitope mRNA/LNP vaccine with antigen-specific immunogenicity and antitumor activity

**DOI:** 10.64898/2026.07.06.736667

**Authors:** Ayushi Verma, Sang Hoon Kim, Ban Seok Lee, Jong Hoo Lee, Hyeryung Kim, Hesung Now, Yoonjoo Choi, Dong-Sup Lee, Woong-Yang Park, Young Ae Park

**Author notes:** Correspondence should be addressed to Y.A. Park.

## Abstract

Personalized neoantigen vaccines are an emerging strategy for cancer immunotherapy, but their effectiveness depends on selecting tumor-specific antigens capable of inducing functional T-cell responses. The VACINUS AI-informed neoantigen prioritization framework previously identified and peptide-validated three immunogenic Tier 1 neoantigens in the B16F10 melanoma model. In this study, we extended that framework by translating these validated neoantigens into a multi-epitope messenger RNA vaccine formulated with lipid nanoparticles and evaluating its preclinical immunogenicity and antitumor activity. The three VACINUS-prioritized B16F10 neoantigens were encoded within a single multi-epitope construct, BF-V1_27-Ser, and formulated to generate BF-RNA-P. In B16F10 tumor-bearing mice, BF-RNA-P induced neoantigen-specific CD8+T-cell responses, with the strongest response directed against the B16F10-1-4 epitope. Combination with anti-PD-1 further enhanced vaccine-induced CD44+IFN-γ+ CD8+ T-cell activation, whereas anti-PD-1 alone did not induce detectable peptide-specific responses. BF-RNA-P also suppressed tumor growth in vivo, and combination treatment produced the strongest antitumor effect, reflected by reduced tumor volume and lower endpoint tumor burden. Together, these findings demonstrate that VACINUS-prioritized and peptide-validated neoantigens can be reformatted into a multi-epitope messenger RNA/lipid nanoparticle vaccine while retaining antigen-specific immunogenicity and antitumor activity. This study provides preclinical proof-of-concept for integrating AI-informed, TCR-aware neoantigen prioritization with messenger RNA/lipid nanoparticle delivery as a translational strategy for personalized cancer vaccine development.

## 1. Introduction

Personalized neoantigen vaccines have emerged as a promising strategy for cancer immunotherapy because they target tumor-specific mutations that are absent from normal tissues. These mutation-derived peptides can be recognized as foreign by the immune system and may therefore induce tumor-specific T-cell responses with limited risk of autoimmunity (1). Clinical studies have shown that neoantigen vaccines can generate both CD8+ and CD4^+^ T-cell responses and contribute to tumor control in several cancer types, including melanoma, glioblastoma, and pancreatic cancer (2–4). However, despite these advances, a major challenge remains:only a small fraction of predicted neoantigens are truly immunogenic.

Conventional neoantigen prediction pipelines commonly prioritize candidates based on somatic mutation calling, tumor expression, and predicted peptide binding to MHC class I molecules (5). Although peptide-MHC binding is required for antigen presentation, it does not guarantee recognition by T cells. In many cases, peptides predicted to bind MHC class I fail to induce measurable T-cell responses after experimental validation (6–8). This gap indicates that antigen presentation alone is not sufficient to identify the most effective vaccine targets.

A critical determinant of neoantigen immunogenicity is T-cell receptor recognition. Productive CD8+ T-cell activation depends on the interaction between the TCR and the peptide-MHC complex, rather than peptide-MHC formation alone (4,5,9). Therefore, incorporating TCR-pMHC recognition into neoantigen prioritization may improve the selection of candidates with true immunogenic potential. To address this, the VACINUS platform was previously developed as an AI -informed neoantigen prediction workflow that integrates peptide-MHC binding prediction with tumor-reactive TIL TCR-pMHC ternary complex modeling. This strategy was designed to prioritize high-confidence Tier 1 neoantigens with an increased likelihood of being recognized by tumor-reactive CD8+ T cells (8).

In the previous VACINUS study, this approach was validated using the B16F10 mouse melanoma model (8). In that study, three Tier 1 neoantigens, B16F10-1-3, B16F10-1-4, and B16F10-1-7, were experimentally confirmed to induce antigen-specific CD8+ T-cell responses. Peptide vaccination with these Tier 1 neoantigens also showed antitumor activity, particularly when combined with anti-PD-1 immune checkpoint blockade. These findings established B16F10-1-3, B16F10-1-4, and B16F10-1-7 as VACINUS-prioritized and peptide-validated immunogenic neoantigens.

Although peptide vaccination is valuable for validating neoantigen immunogenicity, peptide-based platforms have limitations for clinical translation, especially in personalized multi-epitope vaccine development. From an immunological perspective, exogenously administered peptides may have limited in vivo persistence and often depend on efficient uptake and cross-presentation by antigen-presenting cells to induce strong CD8+ T-cell responses. Each peptide usually requires separate synthesis, purification, and formulation, which can increase manufacturing complexity when multiple patient-specific neoantigens are combined (10–12). In contrast, mRNA vaccines provide a flexible and scalable platform in which multiple neoantigen sequences can be encoded within a single construct and rapidly produced by cell-free in vitro transcription (13,14). Importantly, mRNA vaccines enable intracellular production of the encoded antigen in host cells, including antigen-presenting cells, thereby supporting endogenous antigen processing and MHC class I presentation, which is essential for cytotoxic CD8+ T-cell priming (15,16). When formulated with lipid nanoparticles, mRNA vaccines can support efficient cellular uptake, intracellular antigen expression, and antigen presentation through MHC class I pathways, thereby promoting cytotoxic CD8^+^ T-cell priming (17,18). LNP formulation also provides additional immunological advantages by protecting mRNA from degradation, promoting cytosolic delivery, and stimulating innate immune signals that can enhance dendritic-cell activation, antigen presentation, and adaptive T-cell responses (19). These features make the mRNA/LNP platform a potentially adaptable format for translating computationally prioritized and peptide-validated neoantigens into multi-epitope vaccine candidates. Therefore, the rationale for using an mRNA/LNP format in this study was not only manufacturing flexibility, but also its ability to support sustained intracellular antigen expression, MHC class I antigen presentation, innate immune activation, and robust neoantigen-specific CD8+ T-cell priming.

In this study, we performed a preclinical proof-of-concept evaluation to determine whether the immunogenicity of VACINUS-prioritized Tier 1 neoantigens is preserved after conversion from a peptide vaccine format into an mRNA/LNP vaccine. We designed a multi-epitope mRNA construct, BF-V1_27-Ser, encoding the three previously validated B16F10 Tier 1 neoantigens B16F10-1-3, B16F10-1-4, and B16F10-1-7. The BF-V1_27-Ser mRNA was formulated into lipid nanoparticles, and the resulting LNP-formulated mRNA vaccine is referred to as BF-RNA-P. We then evaluated BF-RNA-P through a stepwise translational workflow, including mRNA construct design, LNP physicochemical characterization, antigen-specific CD8+ T-cell responses, antitumor efficacy alone and in combination with anti-PD-1 therapy.

This study was not designed as a formal optimization or head-to-head comparison of cap structure, dose, administration route, lipid composition, or other formulation variables. Rather, its primary objective was to test whether VACINUS-selected, peptide-validated neoantigens could be reformatted into an mRNA/LNP vaccine while retaining measurable antigen-specific immunogenicity and antitumor activity. Together, our findings extend the VACINUS platform from AI-informed neoantigen prioritization and peptide-level validation to mRNA/LNP vaccine development. This work supports the combination of TCR-informed neoantigen prioritization with mRNA/LNP delivery as a translational framework for personalized cancer vaccine development.

## 2. Materials and Methods

### 2.1 Selection of peptide-validated B16F10 neoantigens

The B16F10 murine melanoma model on the C57BL/6 background was used in this study. Peptide-validated neoantigens were selected from the previously published VACINUS neoantigen discovery and prioritization framework described by Kim et al. (8). In the original study, tumor-expressed nonsynonymous somatic mutations were identified using whole-exome sequencing (WES), whole-transcriptome sequencing (WTS), single-cell RNA sequencing, and TCR repertoire analyses of B16F10 tumors, followed by computational neoantigen prioritization and experimental validation (8).

For translational development in the present study, three previously validated Tier 1 neoantigens, B16F10-1-3, B16F10-1-4, and B16F10-1-7, were selected for multi-epitope mRNA construct generation. Each neoantigen sequence included the predicted 9-mer neoepitope core together with adjacent flanking amino acid residues to facilitate intracellular antigen processing and MHC class I presentation. The neoantigen sequences were subsequently arranged in tandem within a single open reading frame to generate a multi-epitope mRNA construct.

### 2.2 Neoantigen prioritization and Tier classification

Details of the VACINUS Algorithm based neoantigen prediction workflow have been previously described by Kim et al. (8). Briefly, all possible 9-mer and 10-mer peptides derived from mutation-containing protein sequences were evaluated for MHC class I binding affinity using MHCflurry (v2.1.1). Predictions were performed for the H2-Kᵇ and H2-Dᵇ alleles corresponding to the C57BL/6 mouse MHC haplotype. Peptides with predicted IC50 values below 140 nM were considered candidate binders. Peptides bearing mutations at the second anchor position (P2) were excluded from further analysis, based on the predefined VACINUS neoantigen prioritization criteria. Candidate peptides were subsequently filtered using the PRIME immunogenicity prediction model (20) with a threshold of 0.847 % rank to enrich potentially immunogenic neoepitopes. To further evaluate tumor-reactive T-cell recognition potential, interactions between peptide–MHC complexes and CDR3β sequences derived from putative tumor-reactive CD8+ tumor-infiltrating lymphocytes (TILs) were analyzed using pMTnet (v1.0) (21). Neoantigens were subsequently classified into Tier 1 or non-Tier 1 groups based on combined peptide-MHC binding and predicted TCR recognition scores, as previously described (8).

### 2.3 EGFP reporter mRNA expression assay for cap-structure comparison

For cap-structure comparison, EGFP-encoding IVT mRNAs were prepared using the same IVT-based synthesis workflow described above. The same EGFP coding sequence was used to generate reporter mRNAs with either ARCA or CleanCap® AG (3′-O-Me) cap analogs. The resulting EGFP reporter mRNAs were subjected to quality assessment by absorbance-based measurement and Tape Station analysis prior to cell-based expression evaluation.

K-562 cells obtained from the Korean Cell Line Bank were maintained in RPMI 1640 medium supplemented with 10% fetal bovine serum and 1% antibiotics-antimycotic solution. EGFP-encoding IVT mRNAs were introduced into K-562 cells by electroporation using a Gene Pulser II apparatus (Bio-Rad). After electroporation, cells recovered in culture medium and incubated for 24 h before flow cytometric analysis. EGFP expression was assessed by flow cytometry using a MACSQuant 10 flow cytometer (Miltenyi Biotec). Live cells were selected based on FSC-A and SSC-A profiles, followed by single-cell gating using FSC-H and FSC-A. EGFP-positive cells were identified from FITC-A histogram profiles, with the gate set using the pulse-only negative control. Mean fluorescence intensity of the EGFP-positive cell population was used as the reporter expression readout. Flow cytometry data were analyzed using FlowJo v10.0.01.

This assay was used as an in vitro reporter-based comparison of ARCA- and CleanCap® AG (3′-O-Me)-capped IVT mRNAs to support selection of the cap structure for subsequent neoantigen-encoding mRNA preparation.

### 2.4 mRNA Synthesis by In Vitro Transcription

#### 2.4.1 Plasmid Construction

The BF-V1_27-Ser coding sequence was cloned into the corresponding Geninus in-house vector backbone for IVT template preparation. Two vector variants were used for cap-specific construct preparation: Geninus Empty Vector A-cap and Geninus Empty Vector C-cap. Both the insert and vector backbone were digested with EcoRI-HF and BamHI-HF restriction enzymes (NEB, R3101S and R3136S), purified using the Axen™ PCR DNA Kit (Macrogen, MG-P-002-200), and ligated using T4 DNA Ligase (Thermo Fisher Scientific, EL0011). The ligation product was introduced into DH5α chemically competent E. coli (Enzynomics, CP011), followed by ampicillin-based colony selection. Selected colonies were cultured, and plasmids were purified using the EZ-Pure™ Plasmid Prep Kit Ver. 2 (Enzynomics, EP101-200N). The presence of the insert and the integrity of the poly(A) tail were assessed by restriction enzyme digestion and band pattern analysis. The insert sequence was confirmed by Sanger sequencing (Macrogen), and no sequence discrepancy was observed compared with the corresponding vector map. The confirmed plasmid constructs, designated Geninus BF-V1_27-Ser A-cap and Geninus BF-V1_27-Ser C-cap, were used for subsequent IVT template preparation.

#### 2.4.2 IVT Template Preparation

The confirmed plasmid construct was used to generate the IVT DNA template by PCR amplification using custom IVT primers (IVT-F and IVT-R) and Phusion High-Fidelity PCR Master Mix with GC Buffer (Thermo Fisher Scientific, F532L). The amplified IVT template was purified using the Axen™ PCR DNA Kit (Macrogen, MG-P-002-200) and subjected to AseI digestion (NEB, R0526S) to remove the primer-derived tail region located outside the poly(A) tail. The resulting IVT DNA template was purified and assessed by absorbance-based dsDNA quantification and TapeStation analysis. DNA concentration and purity were evaluated using NanoDrop, and the expected template size and band pattern were confirmed using a 4200 TapeStation system (Agilent) with D1000 or D5000 Screen Tape, depending on the template length.

#### 2.4.3 mRNA Synthesis and Purification

Capped IVT mRNA was generated using the purified IVT DNA template, T7 RNA polymerase (Enzynomics, RP001L), NTP Set Tris buffered (Thermo Fisher Scientific, R1481), RiboLock RNase Inhibitor (Thermo Fisher Scientific, EO0381), magnesium chloride solution (Sigma-Aldrich, M1028-100ML), and DTT (Thermo Fisher Scientific, P2325). Depending on the experimental design and mRNA batch, either ARCA or CleanCap® AG (3′-O-Me) cap analogs were used. ARCA-capped mRNA was generated using m²⁷,3′-OGP₃G ARCA Cap Analog solution (Jena Bioscience, NU-855L), whereas CleanCap-capped mRNA was generated using CleanCap® Reagent AG (3′-O-Me) (TriLink, N-7413-100). Following IVT, residual DNA template was removed by DNase I treatment using RNase-free DNase I (Thermo Fisher Scientific, EN0521). The resulting capped IVT mRNA was purified by column-based RNA cleanup using the Monarch® Spin RNA Cleanup Kit (NEB, T2050L) prior to LNP formulation.

#### 2.4.4 Geninus Vector Backbones and Functional Elements

Schematic maps of the Geninus in-house Empty Vector A-cap and C-cap backbones and the resulting Geninus BF-V1_27-Ser A-cap and C-cap construct vectors are provided in Supplementary Figure S2. The internal Geninus designations “A-cap” and “C-cap” refer to vector backbone variations and do not correspond to the cap analog reagents (ARCA or CleanCap®) used during in vitro transcription. The vector maps display the major functional elements used for cloning and IVT template preparation, including: (i) T7 promoter for directing RNA polymerase-mediated transcription; (ii) transcription start region defining the beginning of RNA synthesis; (iii) 5′ untranslated region (5′ UTR) regulating mRNA translation efficiency; (iv) multiple cloning site (MCS) or BF-V1_27-Ser coding sequence containing the gene of interest; (v) 3′ untranslated region (3′ UTR) influencing mRNA stability and localization; (vi) poly(A) tail region enhancing mRNA stability and translation; (vii) primer binding sites enabling PCR-based template amplification and IVT primer annealing; (viii) restriction enzyme sites (EcoRI, BamHI, AseI) facilitating directional cloning and template processing; (ix) ampicillin resistance gene enabling antibiotic-based selection of transformed bacteria; and (x) bacterial origin of replication ensuring plasmid replication in E. coli. Full vector sequences are not shown in the supplementary figure but are available upon request.

### 2.5 Lipid nanoparticle (LNP) formulation

Purified IVT mRNA was formulated into lipid nanoparticles (LNPs) using commercially available lipid components purchased from MedChemExpress, including ALC-0315 (HY-138170), ALC-0159 (HY-138300), SM-102 (HY-134541), DMG-PEG 2000 (HY-112764), cholesterol (HY- N0322), and DSPC (HY-W040193). LNPs were prepared using a pipette-based formulation workflow adapted from the manufacturer’s instructions and an internal formulation protocol. Depending on the experimental design, either an ALC-0315-based or an SM-102-based formulation was used. For each formulation batch, purified IVT mRNA and lipid solutions were prepared separately. The mRNA solution was prepared in 100 mM sodium acetate buffer using UltraPure™ DNase/RNase-Free Distilled Water (Invitrogen, 10977023) and sodium acetate buffer, 3 M, pH 5.2 (Sigma-Aldrich, 567422-100ML). Lipid components were dissolved in ethanol (Sigma-Aldrich, 459844-500ML) and combined according to the designated formulation composition. The ALC-0315-based formulation consisted of ALC-0315, DSPC, cholesterol, and ALC-0159 at a molar ratio of 46.3:9.4:42.7:1.6. The SM-102-based formulation consisted of SM-102, DSPC, cholesterol, and DMG-PEG 2000 at a molar ratio of 50:10:38.5:1.5. The mRNA and lipid solutions were combined using a pipette-based mixing procedure to generate mRNA-loaded LNPs. Following formulation, the mRNA-LNP suspension was subjected to stabilization and dialysis-mediated buffer exchange. Samples were dialyzed using Slide-A-Lyzer MINI dialysis devices (10 kDa MWCO, 2 mL; Thermo Fisher Scientific, 88404) against 1× PBS (pH 7.4; Tech & Innovation, BPB-9104). When required, samples were concentrated using Amicon Ultra centrifugal filters (100 kDa MWCO; Millipore, UFC910096). The final formulation was filtered through a 0.22 μm Millex polyether sulfone syringe filter (Millipore, SLGPX13NL). The resulting mRNA-LNP formulations were subjected to quality assessment prior to in vivo administration. For traceability, the corresponding mRNA construct, cap type, LNP formulation type, mRNA batch identifier, and mRNA-LNP batch identifier were recorded for each formulation batch.

### 2.6 Characterization of IVT mRNAs and mRNA-LNP formulations

IVT mRNAs and mRNA-LNP formulations used in this study were analytically characterized prior to experimental use. For IVT mRNAs, RNA concentration and absorbance ratios were measured using a NanoDrop One spectrophotometer (Thermo Fisher Scientific, ND-ONE). A260/A280 and A260/A230 ratios were recorded as absorbance-based sample attributes. RNA size distribution was analyzed using an Agilent 4200 TapeStation system (Agilent Technologies, G2991AA). Briefly, RNA samples were mixed with RNA ScreenTape sample buffer, denatured at 72°C for 3 min, cooled on ice, and analyzed according to the manufacturer’s workflow. The expected RNA profile was confirmed based on the observed main band or peak corresponding to the intended transcript size. For mRNA-LNP formulations, particle size and polydispersity index were measured by dynamic light scattering using a Zeta-potential and particle size analyzer (ELSZ-2000, Otsuka Electronics). Samples were diluted in phosphate-buffered saline and measured at 25°C. mRNA concentration and encapsulation efficiency were determined using the Quant-iT RiboGreen RNA assay kit (Thermo Fisher Scientific). Fluorescence was measured using a Hidex Sense Microplate Reader (Hidex, 425-301) in white/clear 96-well plates (Corning, 3632), after comparison of total RNA and non-encapsulated RNA fractions in the presence or absence of Triton X-100 (Sigma, T8787-100ML), respectively. Batch information and analytical characterization results for the IVT mRNAs and mRNA-LNP formulations are summarized in Supplementary Table S1-A and Supplementary Table S1-B, respectively.

### 2.7 B16F10 cell culture and syngeneic tumor-bearing mouse model

B16F10 murine melanoma cells were obtained from the Korean Cell Line Bank (KCLB, Seoul, Korea; KCLB No. 80008). Cells were thawed from liquid nitrogen storage and cultured in DMEM supplemented with 10% fetal bovine serum at 37°C in a humidified atmosphere containing 5% CO₂. For the main B16F10 tumor-bearing study, male SPF C57BL/6J mice were subcutaneously inoculated in the right flank with 2 × 10⁶ B16F10 cells suspended in 0.1 mL PBS. Mice were randomized into treatment groups when the mean tumor volume reached approximately 60 mm³. The in vivo procedures, including tumor implantation, randomization, test article administration, body-weight monitoring, tumor-volume measurement, survival observation, necropsy, and spleen collection, were performed at the Nonclinical Research Institute, CORESTEMCHEMON Inc., under protocols approved by its Institutional Animal Care and Use Committee. This study was conducted in an AAALAC International-accredited animal facility under IACUC approval no. 2024-0738.

For the dose-dependent immunogenicity assessment shown in Figure 4, a separate B16F10 tumor-bearing mouse experiment was conducted using male SPF C57BL/6NcrlOri mice. In this experiment, B16F10 cells were subcutaneously inoculated in the right flank at 5 × 10⁵ cells in 0.1 mL PBS per mouse, and mice were randomized when the mean tumor volume reached approximately 60 mm³. This experiment was performed at the same external nonclinical research facility under IACUC approval no. 2025-0278 and was analyzed separately from the main B16F10 tumor-bearing study.

### 2.8 mRNA/LNP test articles and treatment regimen

The BF-V1_27-Ser mRNA was used for the in vivo immunogenicity and anti-tumor efficacy studies. For the main B16F10 tumor-bearing study, BF-V1_27-Ser was administered as an LNP-formulated mRNA vaccine. PBS was used as the vehicle control, and empty LNP was used as the lipid carrier control. Anti-mouse PD-1 antibody (BioXCell, BP0146, lot no. 861023M2) was used as an immune checkpoint inhibitor control and as a combination partner with BF-V1_27-Ser mRNA/LNP. In the main tumor-bearing study, mice were assigned to PBS, empty LNP, anti-PD- 1, BF-V1_27-Ser mRNA/LNP, or BF-V1_27-Ser mRNA/LNP plus anti-PD-1 treatment groups. BF-V1_27-Ser mRNA/LNP was administered intravenously at 20 μg per mouse, and anti-mouse PD-1 antibody was administered intraperitoneally at 200 μg per mouse. Treatments were administered once weekly. Mice assigned to immunogenicity analysis received two doses and were sacrificed 7 days after the second treatment for spleen collection and ex vivo analysis, whereas mice assigned to antitumor efficacy assessment received three doses. Additional BF-V1_27-Ser-based mRNA/LNP formulations used in supplementary immunogenicity assessments are described according to the corresponding experimental settings, including mouse strain or condition, dose, route, cap structure, LNP composition, and treatment schedule. Batch-specific information for the mRNA/LNP formulations used in the main tumor-bearing study, the dose-dependent immunogenicity assessment, and the supplementary immunogenicity studies are summarized in Supplementary Table S1 for traceability.

### 2.9 Splenocyte preparation and ex vivo peptide restimulation

Spleens collected for immunogenicity analysis were processed on the day of collection. Each spleen was mechanically dissociated through a 70 μm cell strainer in RPMI 1640 medium supplemented with 1% fetal bovine serum. The cell suspension was centrifuged, and red blood cells were lysed using RBC lysis buffer. After lysis, splenocytes were washed and resuspended in RPMI 1640 medium containing 1% fetal bovine serum. Cell numbers were determined using acridine orange staining and a Countess II automated cell counter. Freshly isolated splenocytes were seeded in 96-well plates at 1 × 10⁶ cells per well and restimulated ex vivo with individual short peptides corresponding to the encoded B16F10 antigen sequences, B16F10-1-3, B16F10-1-4, or B16F10-1-7. These peptides were used only as ex vivo restimulation reagents. Unstimulated cells served as the negative control, and PMA/ionomycin stimulation was used as the positive control. Brefeldin A was added during stimulation to allow intracellular cytokine accumulation. Cells were incubated for 6 h at 37°C in a humidified atmosphere containing 5% CO₂ before antibody staining and flow cytometric analysis.

### 2.10 Flow cytometric analysis of antigen-specific CD8 T cell responses

After ex vivo peptide restimulation, splenocytes were stained for viability and surface T cell markers. The surface staining panel consisted of CD4-Pacific Blue, CD8-FITC, CD44-PE/Cy7, and LIVE/DEAD-APC/Cy7. Antibodies were diluted in FACS buffer and incubated with cells for 30 min at 4°C in the dark. The dilution factors were 1:400 for CD4, CD8, and CD44 antibodies and 1:1,000 for the LIVE/DEAD viability dye. After staining, cells were washed with PBS-based FACS buffer containing 0.5% and 0.1% sodium azide. For intracellular cytokine staining, cells were fixed and permeabilized using a fixation/permeabilization buffer for 10 min. IFN-γ-PE antibody was diluted 1:200 in permeabilization/wash buffer, and cells were incubated for 50 min at 4°C in the dark. After washing, flow cytometry data were acquired using a MACSQuant 10 flow cytometer and analyzed with FlowJo v10.0.01. Lymphocytes were gated based on FSC-A and SSC-A, followed by singlet gating using FSC-H and FSC-W. Viable cells were defined as LIVE/DEAD-negative cells. Within the viable CD8 T cell population, antigen-responsive activated CD8 T cells were quantified as CD44⁺IFN-γ⁺ cells. The frequency of CD44⁺IFN-γ⁺ CD8 T cells was used as the primary readout for antigen-specific cellular immune responses.

### 2.11 Tumor monitoring and efficacy assessment

Tumor volume and body weight were measured twice weekly from the start of treatment. Tumor dimensions were measured using a Vernier caliper along the long and short axes, and tumor volume was calculated using the following formula: tumor volume = long axis × short axis² / 2. Tumor photographs were taken once weekly after grouping, and tumor weight was measured at necropsy. Survival was monitored throughout the study based on daily clinical observation. Moribund or dead animals were weighed and subjected to necropsy. Tumor growth inhibition was calculated relative to the PBS vehicle control group using the following formula: TGI = [1 − tumor volume of treated group / tumor volume of PBS control group] × 100. Survival was analyzed using the Kaplan–Meier method.

### 2.12 Dose-dependent immunogenicity assessment

To evaluate dose-dependent antigen-specific CD8 T cell responses, a separate B16F10 syngeneic tumor-bearing mouse experiment was performed using BF-V1-27-Ser mRNA/LNP. Only the BF-V1-27-Ser treatment groups were included in this analysis. BF-V1-27-Ser mRNA/LNP was administered intravenously at dose levels assigned as 10 or 20 μg per mouse, with adjusted second-dose levels of 7 or 14 μg per mouse, respectively. InVivo Plus anti-mouse PD-1 (CD279) was administered intraperitoneally at 200 μg per mouse where indicated. Mice assigned to the immunogenicity cohort received two weekly doses on Day 1 and Day 8 and were sacrificed on Day 13 for spleen collection. Splenocytes were restimulated ex vivo with the B16F10-1-4 peptide, and antigen-specific CD8 T cell responses were quantified as CD44⁺IFN-γ⁺ cells within the viable CD8 T cell population using the same flow cytometric strategy described above. This dose-dependent immunogenicity experiment was analyzed separately from the main B16F10 tumor-bearing efficacy study.

### 2.13 Statistical analysis

Data are presented as mean ± SD or mean ± SEM, as indicated in the corresponding figure legends. Statistical analyses were performed using GraphPad Prism version 8.0.2 and SPSS Statistics, as appropriate. For comparisons between two groups, an unpaired two-tailed Student’s t-test was used. For comparisons among multiple groups, one-way ANOVA followed by Tukey’s multiple comparisons test was applied. Tumor growth and body-weight changes measured over time were analyzed using a mixed-effects model followed by Tukey’s multiple comparisons test. Survival was analyzed using the Kaplan–Meier method. A P value < 0.05 was considered statistically significant.

## 3. Results

### 3.1 Translational development of VACINUS-prioritized neoantigens into an mRNA/LNP vaccine platform

To establish a translational neoantigen vaccine strategy, we developed an integrated workflow combining AI-assisted neoantigen prioritization (VACINUS algorithm), experimental immunogenicity validation, and mRNA/LNP-based therapeutic delivery (Fig. 1). The present study builds upon the previously established VACINUS framework, which integrates genomic, transcriptomic, single-cell, and TCR repertoire analyses to identify immunogenic neoantigens derived from tumor-expressed nonsynonymous somatic mutations in the B16F10 melanoma model (8). In the original VACINUS study, tumor-derived nonsynonymous somatic mutations identified through whole-exome sequencing (WES) and whole-transcriptome sequencing (WTS) were evaluated using peptide–MHC binding prediction together with tumor-reactive TCR–pMHC interaction modeling to prioritize therapeutically relevant neoantigens. Candidate neoantigens were subsequently classified into Tier 1 and non-Tier 1 groups and experimentally validated through in vivo peptide vaccination assays (8). This workflow enabled identification of multiple Tier 1 neoantigens capable of inducing strong neoantigen-specific CD8+ T-cell responses and antitumor activity in vivo.

**Figure 1.**
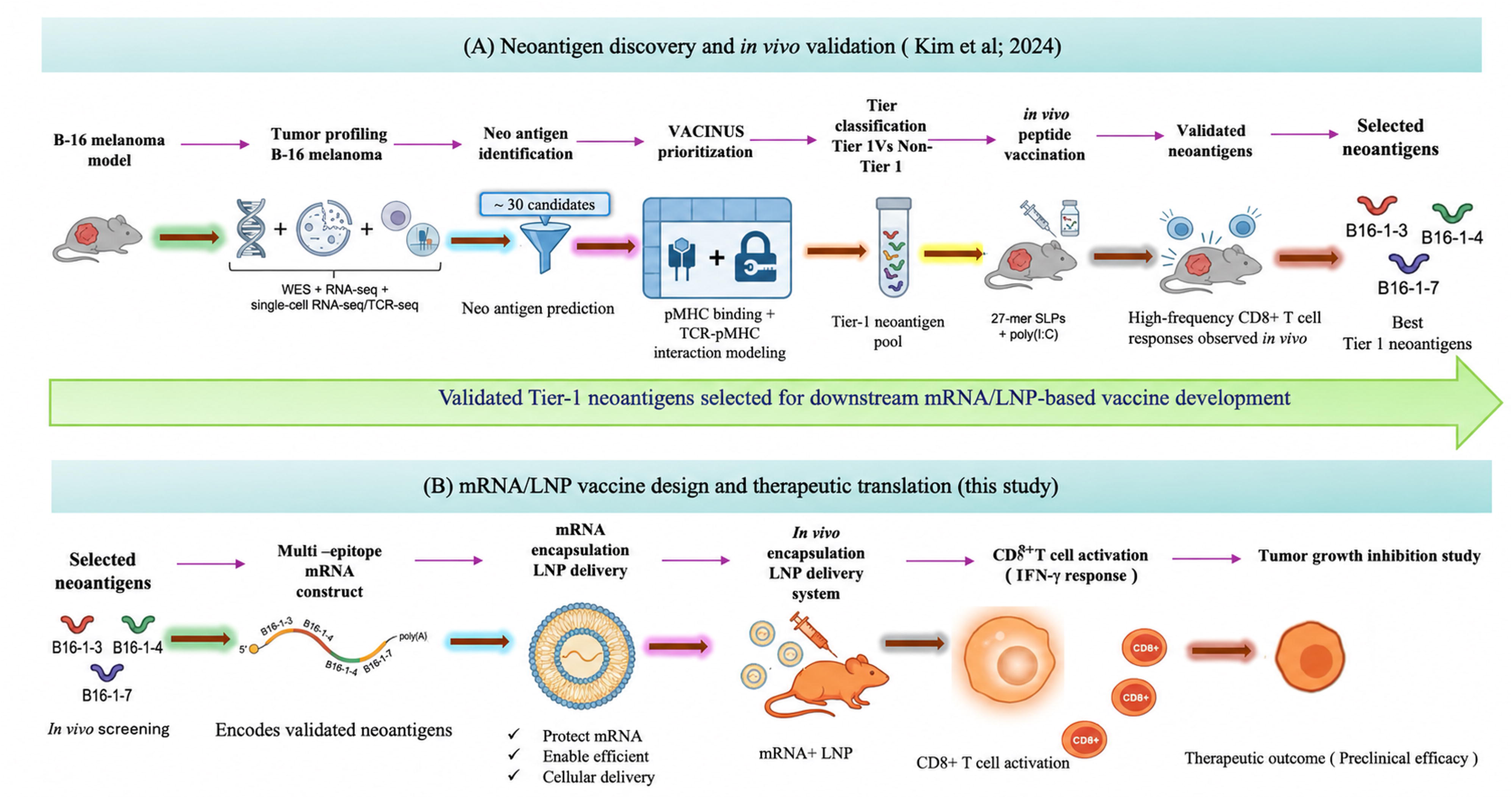
Translational workflow for development of a VACINUS-guided mRNA/LNP neoantigen vaccine platform. (A) Overview of neoantigen discovery and in vivo validation using the previously established VACINUS workflow. Tumor profiling of the B16F10 melanoma model was performed using whole-exome sequencing (WES), RNA-seq, single-cell RNA-seq, and TCR-seq analyses. Candidate neoantigens were prioritized using the VACINUS algorithm, which integrates peptide–MHC binding prediction and tumor-reactive TCR–pMHC interaction modeling. Tier 1 neoantigens were subsequently validated through in vivo peptide vaccination assays, leading to selection of immunogenic neoantigens B16-1-3, B16-1-4, and B16-1-7 showing strong CD8+ T-cell responses. (B) Translational development of validated neoantigens into an mRNA/LNP vaccine platform in the present study. Selected Tier 1 neoantigens were assembled into a multi-epitope mRNA construct and encapsulated into lipid nanoparticles (LNPs) for in vivo delivery. The resulting mRNA/LNP vaccine platform was designed to enable simultaneous expression of multiple validated neoantigens for downstream evaluation of neoantigen-specific immune responses and antitumor therapeutic efficacy. The VACINUS neoantigen prioritization framework was adapted from Kim et al., 2024(8).

Based on these findings, three peptide-validated Tier 1 neoantigens, B16-1-3, B16-1-4, and B16-1-7, were selected for downstream translational development into an mRNA/LNP vaccine platform (Fig. 1A). Unlike the previous peptide-based validation study, the present work focuses on therapeutic translation of VACINUS-prioritized neoantigens into a clinically adaptable mRNA/LNP delivery system for cancer immunotherapy. To generate the vaccine construct, the selected neoantigen sequences were assembled into a multi-epitope mRNA construct containing the predicted neoepitope cores together with flanking amino acid regions to facilitate intracellular antigen processing and MHC class I presentation.

The resulting mRNA construct was subsequently encapsulated into lipid nanoparticles (LNPs) to improve mRNA stability, intracellular delivery, and antigen expression efficiency (Fig. 1B). The resulting mRNA/LNP vaccine platform was designed to enable simultaneous expression of multiple validated neoantigens and promote neoantigen-specific CD8+ T-cell activation, including IFN-γ responses, for downstream evaluation of therapeutic antitumor efficacy in the B16F10 melanoma model. Collectively, this strategy establishes a translational framework linking AI-driven neoantigen discovery and validation with mRNA/LNP-based therapeutic cancer vaccine development.

### 3.2 Designing and Optimizing Multi-Epitope mRNA Constructs from Validated Neoantigens

With three Tier 1 neoantigens confirmed through the VACINUS peptide validation pipeline (Fig. 1A and Supplementary Fig. S1A-C), we moved forward with building a multi-epitope mRNA construct. The selected neoantigens, B16_mv1_7 (B16F10-1-3), B16_mv1_9 (B16F10-1-4), and B16_mv1_8 (B16F10-1-7), were incorporated into a single construct designated BF-V1_27-Ser (Fig. 2), with full sequence details provided in Supplementary Table S2. In BF-V1_27-Ser, the three neoantigen sequences, VSFAPLVQL, CIIRNVQVL, and SAPRASTTA, were arranged in series according to the 7-9-8 configuration within one continuous open reading frame. Rather than inserting artificial linkers between epitopes, we kept the neoantigen-coding regions directly adjacent to preserve the natural antigenic context around each mutation. Each unit retained not just the core neoepitope sequence, but also the flanking amino acid residues surrounding the mutation site. The broader construct architecture was built around human β-globin 5′ and 3′ UTRs, with a segmented poly(A) tail (40+ 60 nt) and cap structures flanking the tandem neoantigen array.

**Figure 2.**
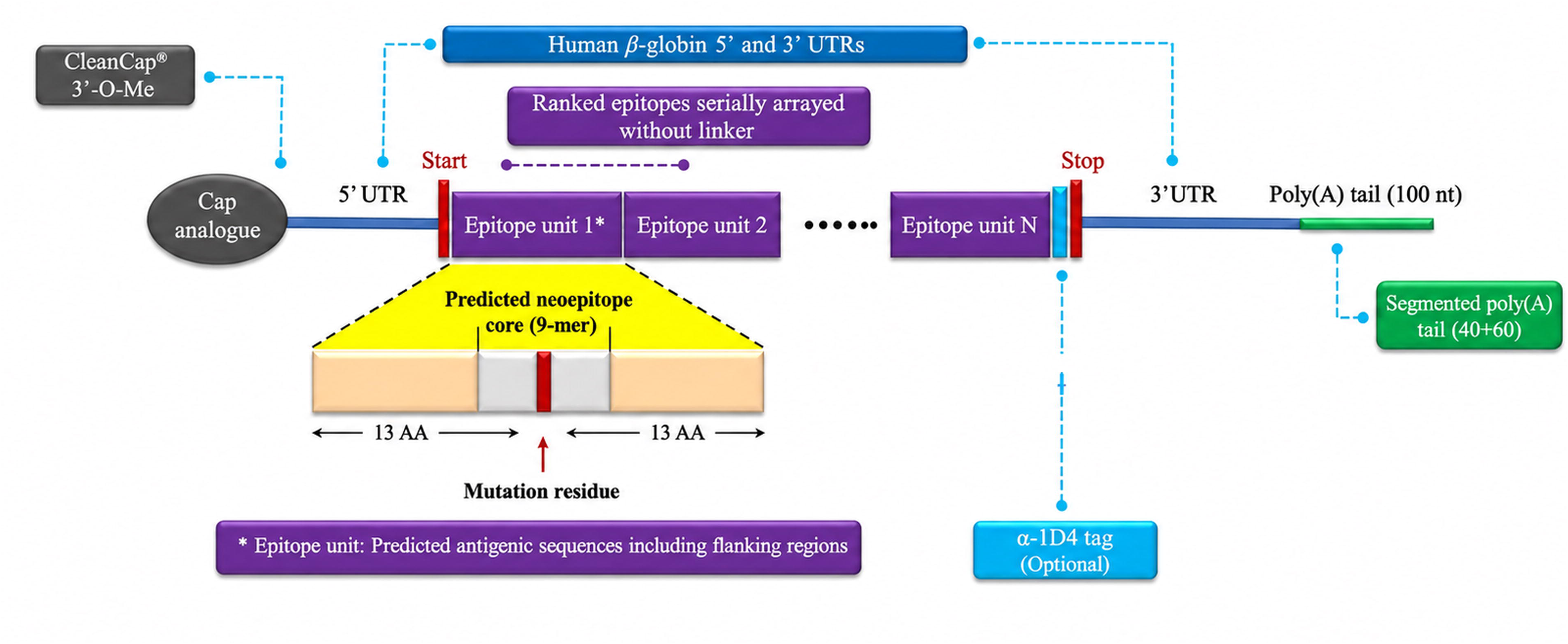
Design and optimization of multi-epitope mRNA constructs encoding validated neoantigens. Schematic overview of the BF-V1_27-Ser construct, built from three Tier 1 neoantigens identified through the VACINUS workflow. The neoantigens were arranged serially within a single ORF, flanked by human β-globin 5′ and 3′ UTRs, and paired with a segmented poly(A) tail and cap structure. An exploratory in silico peptide screening step was also applied to assess potential unintended non-target MHC-I-binding peptides generated from the candidate polyepitope construct regions, based on NetMHCpan-predicted binding affinity below 500 nM for H2-Kᵇ or H2-Dᵇ, with exact intended neoepitope cores and intended-core-overlapping peptides excluded from the final candidate count, as summarized in **Supplementary Table S3.**

During development, we compared two capping strategies, ARCA and CleanCap AG (3′OMe), using an EGFP reporter system. Flow cytometry analysis showed that the CleanCap AG (3′OMe)-capped construct produced substantially higher EGFP expression, with a mean fluorescence intensity (MFI) of 310 compared to 120 for the ARCA-capped construct (Supplementary Fig. 1B, C). These findings suggested that cap structure may influence mRNA expression efficiency under the tested reporter assay condition.

To further evaluate the functional immunogenicity of the two capping strategies, ex vivo peptide stimulation was performed using splenocytes from vaccinated mice. CD44+ IFN-γ+ CD8+ T cell responses were assessed following restimulation with the individual neoepitope peptides B16F10-1-3, B16F10-1-4, and B16F10-1-7. CleanCap-mRNA elicited notably stronger CD8+ T cell responses compared to ARCA-mRNA across the neoepitope conditions, most prominently against B16F10-1-4, with PMA/Ionomycin serving as a positive control (Supplementary Fig. S1D). These results suggest that CleanCap AG (3′-O-Me) may enhance mRNA expression and antigen-specific T cell responses under the tested exploratory conditions. The platform was also designed with scalability in mind, as the same construct framework can be adapted to encode multiple epitopes within a single mRNA, supporting broader multi-neoantigen vaccine configurations.

As an additional exploratory in silico quality check before synthesis and formulation, candidate polyepitope construct regions were screened using VACINUS-derived 8-10-mer peptides and their pre-computed NetMHCpan 4.1 binding affinity values. Peptides with a predicted binding affinity below 500 nM for H2-Kᵇ or H2-Dᵇ were retained as affinity-based candidate MHC-I binders. Candidate peptides corresponding to the exact intended 9-mer neoepitope cores, as well as peptides overlapping with the intended neoepitope cores, were excluded when calculating the final unintended non-target MHC-I-binding peptide candidate count. After applying these predefined exclusions, no final unintended non-target MHC-I-binding peptide candidates were identified in this exploratory screening. These results are summarized in Supplementary Table S3.

### 3.3 Formulation of neoantigen-encoding mRNA into lipid nanoparticles

To enable efficient in vivo delivery of neoantigen-encoding mRNA, purified IVT mRNAs were formulated into lipid nanoparticles (LNPs) using a pipette-based mixing strategy (Fig. 3). The mRNA constructs consisted of a 5′ cap structure, untranslated regions (UTRs), a neoantigen-encoding open reading frame (ORF), and a poly(A) tail (Fig. 3A). Depending on the experimental design, mRNAs were formulated using either ALC-0315- or SM-102-based lipid compositions. The formulation workflow involved preparation of mRNA and lipid solutions, mixing to generate mRNA-LNP complexes, stabilization, dialysis-mediated buffer exchange, concentration when required, sterile filtration, and quality assessment prior to in vivo administration (Fig. 3B). The resulting LNPs consisted of ionizable lipids, phospholipids, cholesterol, and PEG-lipids surrounding the encapsulated mRNA cargo (Fig. 3C). Following cellular uptake by endocytosis, protonation of ionizable lipids under acidic endosomal conditions facilitates endosomal escape and cytoplasmic release of mRNA, enabling antigen expression and subsequent MHC-I-mediated antigen presentation (Fig. 3D). This formulation platform was used for all subsequent mRNA-LNP vaccine studies described in this work.

**Figure 3.**
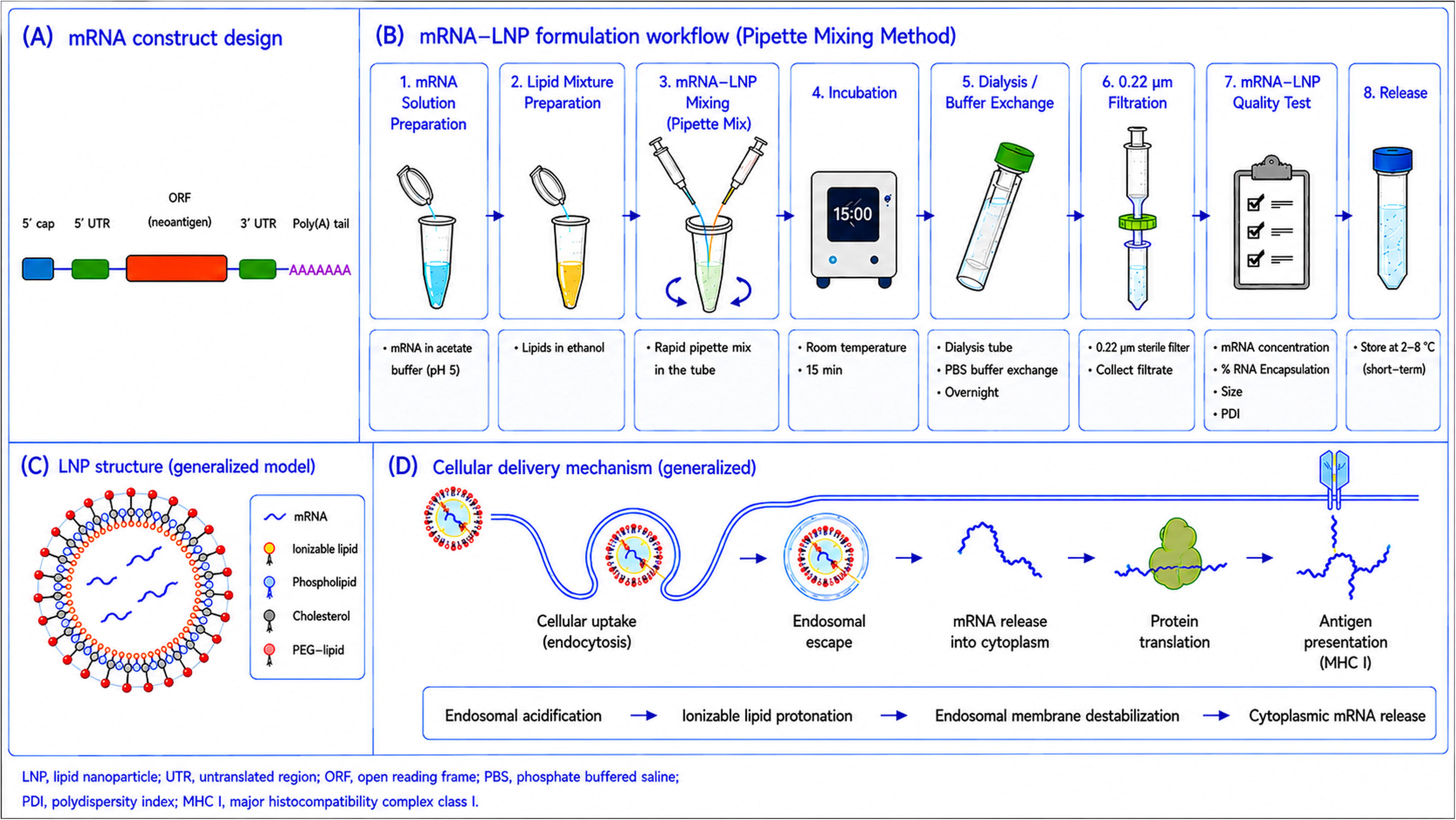
Formulation and delivery mechanism of neoantigen-encoding mRNA-loaded lipid nanoparticles. (A) Schematic representation of the neoantigen-encoding mRNA construct comprising a 5′ cap, 5′ untranslated region (UTR), neoantigen open reading frame (ORF), 3′ UTR, and poly(A) tail. (B) Workflow for mRNA-LNP formulation. Purified IVT mRNA prepared in sodium acetate buffer was mixed with lipid components dissolved in ethanol, followed by stabilization, dialysis-mediated buffer exchange, concentration when required, sterile filtration through a 0.22 μm filter, and quality assessment prior to use. (C) Schematic illustration of the mRNA-loaded lipid nanoparticle structure showing encapsulated mRNA surrounded by ionizable lipids, phospholipids, cholesterol, and PEG-lipids. (D) Proposed intracellular delivery mechanism of mRNA-LNPs. Following endocytic uptake, endosomal acidification promotes protonation of ionizable lipids, leading to membrane destabilization and endosomal escape. Released mRNA is translated in the cytoplasm, and the resulting antigen is processed and presented through the major histocompatibility complex class I (MHC-I) pathway. Abbreviations: LNP, lipid nanoparticle; UTR, untranslated region; ORF, open reading frame; PEG, polyethylene glycol; MHC-I, major histocompatibility complex class I.

### 3.4 BF-V1_27-Ser vaccination elicits B16F10-1-4-specific CD8+ T-cell responses that are enhanced by anti-PD-1 treatment

To evaluate whether BF-V1_27-Ser mRNA/LNP vaccination induces neoantigen-specific cellular immunity, syngeneic B16F10 tumor-bearing C57BL/6J mice were assigned to one of four treatment groups: vehicle control (G1), anti-PD-1 (G2), BF-V1_27-Ser mRNA/LNP (G3), or BF-V1_27-Ser mRNA/LNP plus anti-PD-1 (G4). The experimental workflow is illustrated in Figure 4A. Mice received treatments on Days 0 and 7, and spleens were harvested on Day 14. Splenocytes were isolated and re-stimulated ex vivo with individual B16F10 neoantigen peptides (B16F10-1-3, B16F10-1-4, and B16F10-1-7) to assess antigen-specific recall responses.

**Figure 4.**
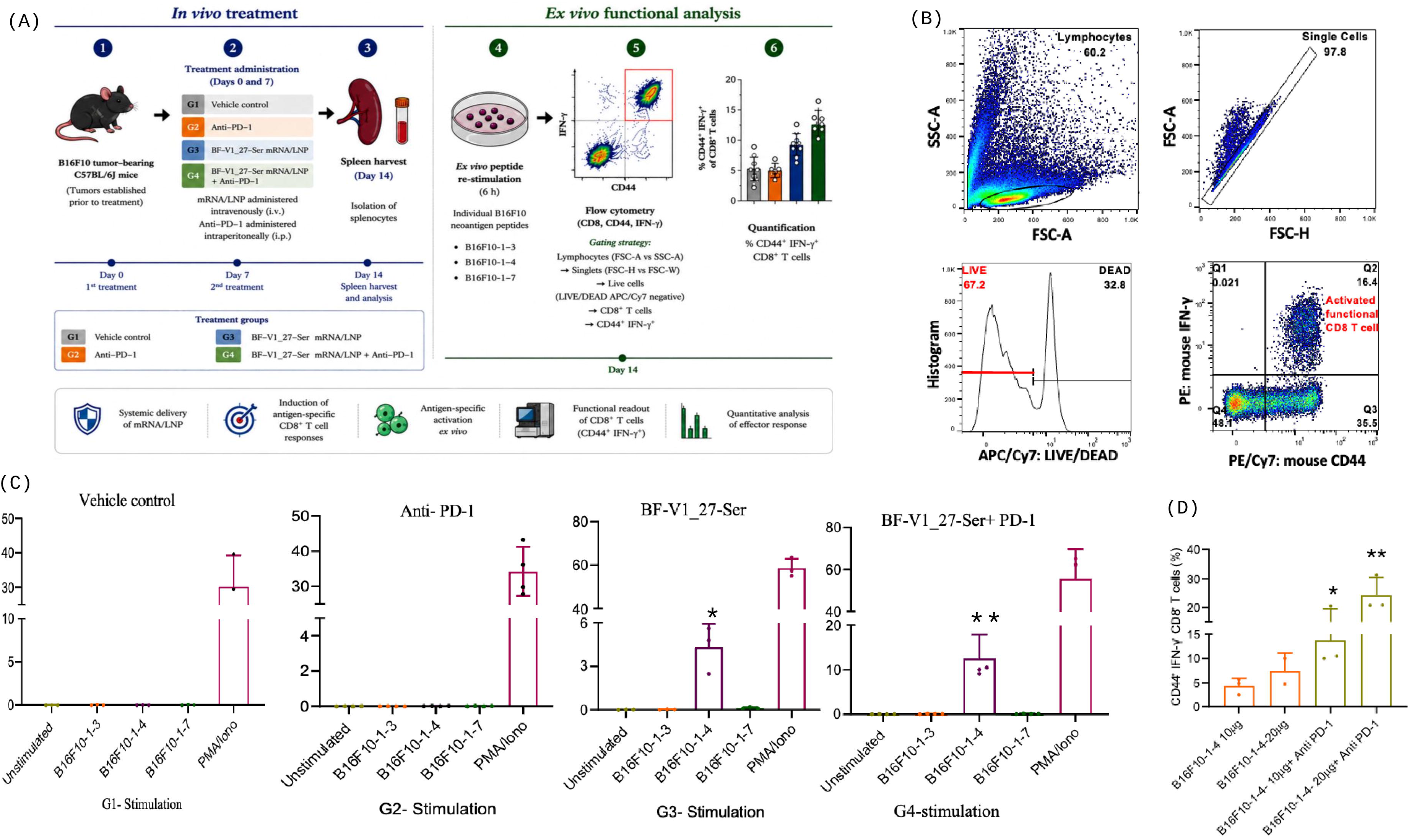
BF-V1_27-Ser vaccination induces neoantigen-specific CD8+ T-cell responses that are enhanced by anti-PD-1 treatment. (A) Schematic overview of the experimental workflow. Syngeneic B16F10 tumor-bearing C57BL/6J mice were assigned to one of four treatment groups: vehicle control (G1), anti-PD-1 (G2), BF-V1_27-Ser mRNA/LNP (G3), or BF-V1_27-Ser mRNA/LNP plus anti-PD-1 (G4). Treatments were administered on Days 0 and 7. Spleens were harvested on Day 14, and splenocytes were isolated for ex vivo functional analysis. Isolated splenocytes were subsequently re-stimulated with individual B16F10-derived neoantigen peptides (B16F10-1-3, B16F10-1-4, and B16F10-1-7) prior to flow cytometric analysis. (B) Flow cytometric gating strategy used to identify activated antigen-specific CD8+ T cells following peptide re-stimulation. Lymphocytes were first gated based on FSC-A and SSC-A characteristics, followed by singlet discrimination using FSC-A and FSC-H parameters. Dead cells were excluded using LIVE/DEAD APC/Cy7 staining. Viable CD8+ T cells were subsequently analyzed for CD44 and intracellular IFN-γ expression. Antigen-specific responses were quantified as the frequency of CD44+IFN-γ+ cells within the viable CD8+ T-cell population. (C) Antigen-specific CD8+ T-cell responses following ex vivo peptide re-stimulation. Splenocytes isolated from vehicle-treated (G1), anti-PD-1-treated (G2), BF-V1_27-Ser-vaccinated (G3), or BF-V1_27-Ser plus anti-PD-1-treated (G4) mice were re-stimulated with B16F10-1-3, B16F10-1-4, or B16F10-1-7 peptides. The frequency of CD44⁺IFN-γ⁺ CD8+ T cells was determined by intracellular cytokine staining and flow cytometry. BF-V1_27-Ser vaccination induced a significant B16F10-1-4-specific CD8+ T-cell response (*p* = *0.0103*, unpaired two-tailed Student’s t-test), which was further enhanced by anti-PD-1 combination treatment (*p value = 0.0034,* unpaired two-tailed Student’s t-test). No detectable peptide-specific responses were observed in the vehicle or anti-PD-1 monotherapy groups or following re-stimulation with B16F10-1-3 and B16F10-1-7 peptides. (D) Quantification of B16F10-1-4-specific CD8+ T-cell responses following ex vivo peptide re-stimulation. Splenocytes isolated from mice treated with 10 μg or 20 μg BF-V1_27-Ser, with or without anti-PD-1 treatment, were re-stimulated with the B16F10-1-4 peptide and analyzed for the frequency of CD44+IFN-γ⁺+CD8+ T cells. Anti-PD-1 combination treatment enhanced vaccine-induced antigen-specific responses, with the highest response observed in the 20 μg BF-V1_27-Ser plus anti-PD-1 group. Panel D was generated from an independent B16F10 tumor-bearing mouse experiment designed to assess dose-dependent immunogenicity and was analyzed separately from the experiment shown in panel C. Data are presented as mean ± SD. Each dot represents an independent flow cytometric analysis of ex vivo peptide-restimulated splenocyte samples. The number of replicates for each group is represented by the number of data points shown. Statistical significance was determined using an unpaired two-tailed Student’s t-test for panel C and Tukey’s multiple comparisons test for panel D. Exact P values are indicated in the figure. *P < 0.05; **P < 0.01; ns, not significant.

Antigen-specific CD8+ T-cell activation was assessed by intracellular cytokine staining and flow cytometry. As shown in Figure 4B, lymphocytes were first gated based on FSC-A and SSC-A characteristics, followed by singlet selection and exclusion of dead cells using LIVE/DEAD APC/Cy7 staining. Viable CD8+ T cells were subsequently analyzed for CD44 and intracellular IFN-γ expression. Antigen-specific responses were quantified as the frequency of CD44⁺IFN-γ⁺ cells within the viable CD8+ T-cell population. The resulting antigen-specific CD8+ T-cell responses are shown in Figure 4C.

Splenocytes isolated from vehicle-treated mice (G1) and anti-PD-1-treated mice (G2) did not show a meaningful increase in CD44+IFN-γ+ CD8+ T-cell responses following re-stimulation with B16F10-1-3, B16F10-1-4, or B16F10-1-7 peptides. In contrast, splenocytes from BF-V1_27-Ser-vaccinated mice (G3 stimulation) exhibited a significant increase in CD44+IFN-γ+CD8+ T cells specifically following re-stimulation with the B16F10-1-4 peptide (*p value = 0.0103)*, whereas responses to B16F10-1-3 and B16F10-1-7 remained low. These findings identify B16F10-1-4 as the dominant responder among the three tested B16F10 neoantigen peptides within the BF-V1_27-Ser vaccine construct. Notably, splenocytes from mice receiving BF-V1_27-Ser in combination with anti-PD-1 (G4 stimulation) displayed a further increase in B16F10-1-4-specific CD44+IFN-γ+ CD8+ T-cell responses (*P = 0.0034)* indicating that PD-1 blockade enhances vaccine-induced antigen-specific CD8+ T-cell activation. Consistent with the known mechanism of action of immune checkpoint blockade, anti-PD-1 treatment alone did not show a meaningful increase in peptide-specific responses, suggesting that PD-1 inhibition amplifies vaccine-primed T-cell responses rather than initiating de novo neoantigen-specific immunity. To further investigate the contribution of vaccine dose and checkpoint blockade, splenocytes from mice immunized with either 10 μg or 20 μg BF-V1_27-Ser, with or without anti-PD-1 treatment, were re-stimulated with the immunodominant B16F10-1-4 peptide (Figure 4D).

Although no significant difference was observed between the 10 μg and 20 μg BF-V1_27-Ser monotherapy groups, anti-PD-1 combination treatment increased the frequency of B16F10-1-4-responsive CD44⁺IFN-γ⁺ CD8+ T cells. The highest response was observed in the 20 μg BF-V1_27-Ser plus anti-PD-1 group, which showed significantly greater T-cell activation compared with the 10 μg BF-V1_27-Ser group (*P = 0.0059*) and the 20 μg BF-V1_27-Ser group (*P = 0.0248*). These findings further support the ability of PD-1 blockade to potentiate vaccine-induced neoantigen-specific cellular immunity.

Altogether, these results suggest that BF-V1_27-Ser mRNA/LNP vaccination induces neoantigen-specific CD8+ T-cell responses predominantly directed against the B16F10-1-4 epitope, and that anti-PD-1 treatment further enhances the magnitude of this vaccine-induced cellular immune response.

### 3.5 BF-RNA-P combined with anti-PD-1 improves antitumor efficacy in B16F10 melanoma-bearing mice

To evaluate the antitumor efficacy of the LNP-formulated BF-V1_27-Ser mRNA vaccine, hereafter referred to as BF-RNA-P, either alone or in combination with immune checkpoint blockade, B16F10 melanoma-bearing C57BL/6J mice were treated according to the experimental schedule shown in Fig. 5A. Mice were initially assigned to five groups with n = 12 mice per group: vehicle control (G1), LNP control (G2), anti-PD-1 monotherapy (G3), BF-RNA-P monotherapy (G4), and BF-RNA-P plus anti-PD-1 combination therapy (G5). BF-RNA-P was administered intravenously, whereas anti-PD-1 antibody was administered intraperitoneally. Tumor growth, body weight, endpoint tumor weight, and survival were monitored throughout the study.

**Figure 5.**
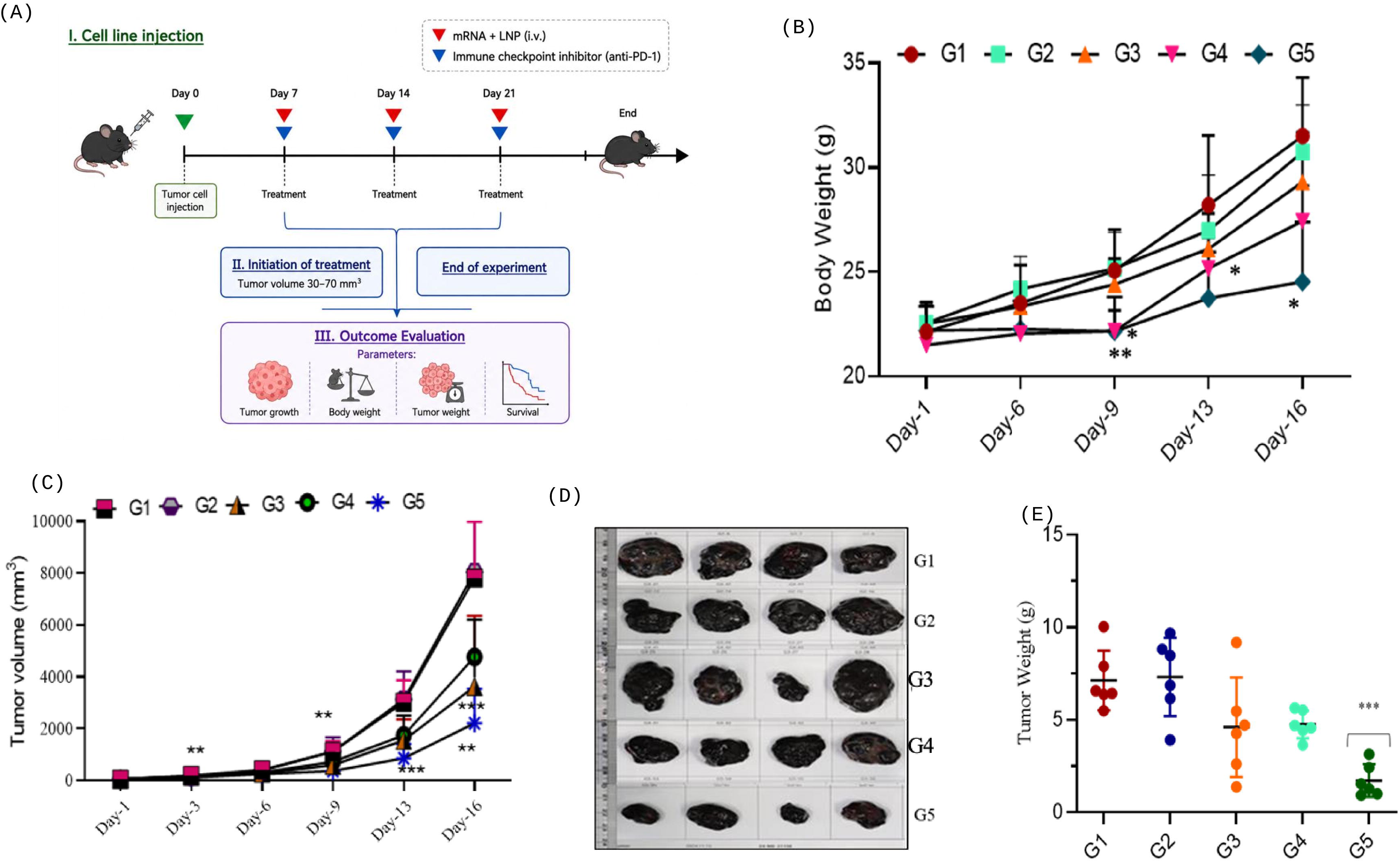
Antitumor efficacy of BF-RNA-P mRNA/LNP vaccination alone or in combination with anti-PD-1 in B16F10 melanoma-bearing mice. (A) Experimental design of the in vivo antitumor efficacy study. B16F10 melanoma cells were inoculated into C57BL/6J mice on Day 0. After tumor establishment, mice were treated with vehicle control, LNP control, anti-PD-1 antibody, BF-RNA-P, or BF-RNA-P plus anti-PD-1 according to the indicated schedule. BF-RNA-P was administered intravenously at 20 µg/head/time, and anti-PD-1 antibody was administered intraperitoneally at 200 µg/head/time. Tumor volume, body weight, endpoint tumor weight, and survival were monitored during the experimental period. (B) Body weight changes during the treatment period. Body weight was measured on Days 1, 6, 9, 13, and 16. Data are presented as mean ± SEM. Statistical significance was determined using a mixed-effects model followed by Tukey’s multiple comparisons test. Significant differences were observed for G4 and G5 at later time points. On Day 9, G4 was significantly lower than G1, G2, and G3 (adjusted P = 0.0079, P = 0.0025, and P = 0.0007, respectively), and G5 was significantly lower than G1, G2, and G3 (adjusted P = 0.0111, P = 0.0126, and P = 0.0071, respectively). On Day 13, G5 was significantly lower than G1 (adjusted P = 0.0310). On Day 16, G5 was significantly lower than G1, G2, and G3 (adjusted P = 0.0195, P = 0.0211, and P = 0.0498, respectively), while G4 was significantly lower than G2 (adjusted P = 0.0403). (C) Tumor growth curves following treatment. Tumor volume was measured on Days 1, 3, 6, 9, 13, and 16 using calipers and calculated as: tumor volume = long axis × short axis² / 2. Data are presented as mean ± SEM. Statistical significance was determined using a mixed-effects model followed by Tukey’s multiple comparisons test. LNP alone did not significantly inhibit tumor growth compared with vehicle control. On Day 3, G5 showed significantly reduced tumor volume compared with G1, G2, and G3 (adjusted *P* = 0.0039, *P* = 0.0113, and *P* = 0.0052, respectively). On Day 9, G3 was significantly lower than G1 and G2 (adjusted P = 0.0146 and P = 0.0255, respectively), while G5 was significantly lower than G1, G2, and G4 (adjusted P = 0.0030, *P* = 0.0018, and *P* = 0.0013, respectively). On Day 13, G3, G4, and G5 were significantly lower than G1 and/or G2, with the strongest effect in G5 (G1 vs. G5, adjusted P = 0.0004; G2 vs. G5, adjusted P < 0.0001). G5 was also significantly lower than G4 on Day 13 (adjusted P = 0.0295). On Day 16, G5 remained significantly lower than G1 and G2 (adjusted P = 0.0098 and P = 0.0034, respectively), and G3 was significantly lower than G2 (adjusted P = 0.0204). (D) Representative images of excised tumors collected at the experimental endpoint from each treatment group. Tumors from mice treated with BF-RNA-P plus anti-PD-1 were visibly smaller than those from the vehicle control, LNP control, and monotherapy groups. (E) Endpoint tumor weight. Each dot represents an individual mouse, and data are presented as mean ± SEM. Tumor weight was analyzed by one-way ANOVA, which showed a significant overall difference among groups (P < 0.0001). The BF-RNA-P plus anti-PD-1 group showed the lowest endpoint tumor weight. Treatment groups: G1, vehicle control (PBS, i.v.); G2, LNP control (i.v.); G3, anti-PD-1 antibody (200 µg/head/time, i.p.); G4, BF-RNA-P (20 µg/head/time, i.v.); G5, anti-PD-1 antibody plus BF-RNA-P. Mice were initially assigned at n = 12 per group. Due to scheduled necropsy and mortality during the study, the number of animals analyzed varied at later time points and endpoint measurements. Statistical significance: ns, not significant; *P < 0.05; **P < 0.01; ***P < 0.001; ****P < 0.0001.

Body weight was measured during the experimental period as a supportive tolerability-related observation (Fig. 5B). No significant difference in body weight was observed among the groups on Day 1. At later time points, body weight was lower in mice receiving BF-RNA-P-containing treatments. On Day 9, BF-RNA-P monotherapy (G4) showed significantly lower body weight compared with G1, G2, and G3 (G1 vs. G4, adjusted *P =* 0.0079; G2 vs. G4, adjusted *P* = 0.0025; G3 vs. G4, adjusted *P* = 0.0007). Similarly, the BF-RNA-P plus anti-PD-1 group (G5) showed significantly lower body weight compared with G1, G2, and G3 on Day 9 (G1 vs. G5, adjusted *P* = 0.0111; G2 vs. G5, adjusted *P* = 0.0126; G3 vs. G5, adjusted *P* = 0.0071). G5 remained significantly lower than G1 on Day 13 (adjusted *P* = 0.0310) and Day 16 (adjusted *P* = 0.0195). On Day 16, G5 was also significantly lower than G2 and G3 (G2 vs. G5, adjusted *P* = 0.0211; G3 vs. G5, adjusted *P* = 0.0498). No significant difference in body weight was observed between G4 and G5 at any time point. No abnormal clinical signs were observed during the experimental period, and survival was not significantly different among treatment groups compared with vehicle control. Tumor growth analysis demonstrated rapid tumor progression in the vehicle control (G1) and LNP control (G2) groups (Fig. 5C). LNP alone did not significantly reduce tumor volume compared with vehicle control at any time point, suggesting that the carrier alone did not contribute appreciably to tumor growth inhibition under these conditions. Anti-PD-1 monotherapy (G3) significantly reduced tumor volume compared with G1 on Day 9 and Day 13 (adjusted *P =* 0.0146 and *P =* 0.0162, respectively), and compared with G2 on Days 9, 13, and 16 (adjusted *P* = 0.0255, *P* = 0.0205, and *P* = 0.0204, respectively). BF-RNA-P monotherapy (G4) also suppressed tumor growth, with significant reductions compared with G1 and G2 on Day 13 (G1 vs. G4, adjusted P = 0.0067; G2 vs. G4, adjusted P = 0.0280).

Notably, BF-RNA-P combined with anti-PD-1 (G5) produced the strongest inhibition of tumor growth. Tumor volume in G5 was significantly reduced compared with G1 on Days 3, 9, 13, and 16 (adjusted *P* = 0.0039, *P* = 0.0030, *P* = 0.0004, and *P* = 0.0098, respectively). G5 was also significantly reduced compared with G2 on Days 3, 9, 13, and 16 (adjusted *P* = 0.0113, *P* = 0.0018, *P* value < 0.0001, and *P* = 0.0034, respectively). Importantly, the combination treatment significantly reduced tumor volume compared with BF-RNA-P monotherapy on Day 9 and Day 13 (G4 vs. G5, adjusted *P* = 0.0013 and *P* = 0.0295, respectively), indicating that anti-PD-1 further enhanced the antitumor effect of BF-RNA-P vaccination. By Day 16, G5 maintained the lowest tumor volume among all groups, although the difference between G4 and G5 did not reach statistical significance at this time point (adjusted *P* = 0.0728).

Consistent with the tumor growth curves, representative images of excised tumors showed visibly smaller tumors in the BF-RNA-P plus anti-PD-1 group compared with the vehicle, LNP, and monotherapy groups (Fig. 5D). Endpoint tumor weight analysis further supported the enhanced antitumor activity of the combination treatment (Fig. 5E). One-way ANOVA showed a significant overall difference in tumor weight among groups (P value < 0.0001), with the BF-RNA-P plus anti-PD-1 group showing the lowest endpoint tumor burden. Together, these results suggest that BF-RNA-P suppresses B16F10 melanoma growth in vivo and that combination with anti-PD-1 further enhances its antitumor activity under the tested conditions.

## 4. Discussion

In this study, we developed a multi-epitope mRNA/LNP vaccine encoding VACINUS-prioritized and peptide-validated neoantigens from the B16F10 melanoma model. The central aim was to determine whether neoantigens selected by the VACINUS-AI informed neoantigen prioritization algorithm and previously validated in peptide-vaccine experiments could be reformatted into an mRNA/LNP vaccine while retaining measurable immunogenicity and antitumor activity(8). The BF-V1_27-Ser mRNA construct, encoding three validated B16F10 neoantigens, was successfully formulated into LNPs, and the resulting vaccine formulation, referred to as BF-RNA-P, induced antigen-specific CD8+T-cell responses and antitumor activity in vivo.

This work extends the previous VACINUS study, in which tumor-derived neoantigens were prioritized using an AI- informed framework integrating peptide-MHC binding prediction with tumor-reactive TCR-pMHC interaction modeling(8). Since MHC binding alone does not reliably predict immunogenicity, incorporating TCR-recognition potential may improve the selection of neoantigens with true functional relevance (6,7,9,22,23). Thus, the novelty of the present study lies not in rediscovering B16F10 neoantigens, but in translating VACINUS-prioritized and peptide-validated targets into a multi-epitope mRNA/LNP vaccine format.

The multi-epitope design of BF-V1_27-Ser is relevant for personalized cancer vaccine development, where simultaneous delivery of multiple patient-specific neoantigens is often required. mRNA provides a flexible platform in which multiple antigenic sequences can be encoded within a single construct and rapidly produced by in vitro transcription (13,14).Compared with peptide-based approaches, mRNA vaccines may simplify multi-epitope vaccine manufacturing and enable intracellular antigen expression, endogenous antigen processing, and MHC class I presentation (10–12,17,18). In this study, the three selected B16F10 neoantigens were incorporated into one mRNA cassette together with flanking amino acid sequences, providing a practical framework for converting validated neoantigen peptides into an mRNA-encoded vaccine. The immunogenicity data indicate that BF-RNA-P preserved antigen-specific T-cell activity after conversion from peptide to mRNA format. Among the three encoded neoantigens, B16F10-1-4 emerged as the dominant immunogenic epitope, inducing a significant CD44⁺IFN-γ⁺ CD8+ T-cell response after ex vivo peptide restimulation. In contrast, responses to B16F10-1-3 and B16F10-1-7 were low under the tested conditions. This epitope hierarchy is biologically plausible, as immunodominance can be shaped by antigen processing, MHC binding, peptide abundance, TCR repertoire availability, and competition among epitopes within a multi-antigen construct(24–26). Therefore, the unequal responses across encoded neoantigens should not be interpreted as failure of the platform, but rather as evidence that individual epitope immunogenicity must still be experimentally evaluated after mRNA construct design.

Combination with anti-PD-1 further enhanced vaccine-associated immune responses. Anti-PD-1 alone did not show a meaningful increase in peptide-specific CD8+ T-cell responses, whereas BF-RNA-P plus anti-PD-1 increased the B16F10-1-4-specific CD44+IFN-γ+ CD8+ T-cell response. This suggests that PD-1 blockade may have amplified vaccine-primed T-cell responses rather than initiating a strong de novo neoantigen-specific response under the tested conditions. This interpretation is consistent with the established role of immune checkpoint blockade in enhancing pre-existing antitumor T-cell responses (24,25,27). These findings support the rationale for combining personalized neoantigen vaccines with immune checkpoint blockade, particularly when vaccination generates antigen-specific T cells that can be further strengthened by release from inhibitory signaling. Similar combination concepts have been supported by clinical and preclinical neoantigen vaccine studies showing that neoantigen vaccination can induce tumor-specific T-cell responses and may cooperate with checkpoint inhibition.

The antitumor efficacy data further support the therapeutic relevance of BF-RNA-P. BF-RNA-P monotherapy suppressed B16F10 tumor growth compared with vehicle and LNP controls, suggesting that the observed antitumor effect was not attributable to the carrier alone. Notably, BF-RNA-P combined with anti-PD-1 produced the strongest tumor-growth inhibition and the lowest endpoint tumor burden. Together with the immunogenicity data, these findings suggest that BF-RNA-P-induced antigen-specific cellular immune responses may contribute to the observed antitumor activity, and that anti-PD-1 may further enhance this vaccine-associated response in the tumor-bearing host. However, because tumor-infiltrating T cells and causal effector mechanisms were not directly dissected in this study, this interpretation should be considered supportive rather than definitive. Similar strategies have been explored in personalized neoantigen vaccine studies, where checkpoint blockade may strengthen vaccine-induced antitumor immunity (2,4).

The formulation-related findings should be interpreted within the exploratory scope of this study. In exploratory experiments, cap structure appeared to influence reporter expression and antigen-specific immune readouts, with CleanCap AG-capped mRNA showing higher values than ARCA-capped mRNA under the tested conditions. However, the study was not designed as a fully matched head-to-head optimization of cap chemistry, dose, lipid composition, or administration route. Because some assays differed in formulation settings and experimental conditions, these data should be presented as exploratory formulation-development observations rather than definitive evidence that CleanCap, a specific dose, lipid composition, or route was formally optimized for the entire study.

Several limitations should be considered. First, this was a preclinical proof-of-concept study rather than a fully matched formulation-optimization or mechanism-dissection study. Second, although antigen-specific IFN-γ-producing CD8+ T cells were detected, the breadth, phenotype, memory potential, clonality, and tumor infiltration of vaccine-induced T cells were not fully characterized. Third, the study focused on the B16F10 melanoma model and three previously validated VACINUS-prioritized neoantigens; therefore, additional tumor models and patient-derived neoantigen sets will be needed to assess broader applicability.

In conclusion, this study extends the VACINUS platform from AI-informed neoantigen prioritization and peptide-level validation to mRNA/LNP-based vaccine development. BF-RNA-P vaccination induced measurable neoantigen-specific CD8+T-cell responses, showed antitumor activity in the B16F10 melanoma model, and demonstrated enhanced efficacy when combined with anti-PD-1 therapy. Importantly, these findings should be interpreted as preclinical proof-of-concept evidence rather than as a definitive comparison of cap structure, dose, administration route, lipid composition, or formulation variables. Future studies incorporating matched formulation comparisons, deeper immune profiling, mechanistic cytotoxicity assays, and additional tumor models will be important to refine this platform and evaluate its broader translational potential for personalized cancer immunotherapy.

## Data Availability Statement

The data supporting the findings of this study are included in the article and its Supplementary Material. Additional data supporting the conclusions of this article will be made available by the corresponding author upon reasonable request, without undue reservation.

## Ethics Statement

No new research on human subjects was conducted in this study. The animal study was reviewed and approved by the Institutional Animal Care and Use Committee of the Nonclinical Research Institute, CorestemChemon Inc. The experiments were conducted in an AAALAC International-accredited animal facility under IACUC approval nos. 2024-0738 and 2025-0278.

## Author Contributions

AV: Formal analysis, Visualization, Writing - original draft, Writing - review and editing. SHK: Methodology, Investigation, Data curation. BSL: Methodology, Investigation. JHL: Software, Formal analysis, Methodology, Data curation. HK: Methodology, Formal analysis, Data curation. HN: Supervision, writing - review and editing. YC: Methodology, Formal analysis, Supervision. DSL: Conceptualization, Methodology, Formal analysis, Supervision. WYP: Conceptualization, Supervision, Funding acquisition, Writing - review and editing. YAP: Conceptualization, Methodology, Formal analysis, Visualization, Writing - original draft, Writing - review & editing, Project administration, Supervision. All authors contributed to the article and approved the submitted version.

## Funding

This research was supported by the Korea Drug Development Fund (KDDF), funded by the Ministry of Science and ICT, the Ministry of Trade, Industry and Energy, and the Ministry of Health and Welfare (Project Unique No. 2710092968, Institutional Research and Development Project No. RS-2023-00284167, Republic of Korea).

## Acknowledgement

The authors gratefully acknowledge Nonclinical Research Institute, CORESTEMCHEMON Inc. for performing the dosing procedures and supplying the biological samples; all animal procedures were IACUC-approved and carried out at an AAALAC-accredited facility.

## Conflict of Interest

Authors affiliated with Geninus Inc. are employees of Geninus Inc., which develops the VACINUS platform. The authors declare that the research was conducted in the absence of any commercial or financial relationships that could be construed as a potential conflict of interest.

