## Supplementary Figures for "AI-Informed neoantigen prioritization enables a multi-epitope mRNA/LNP vaccine with antigen-specific immunogenicity and antitumor activity"

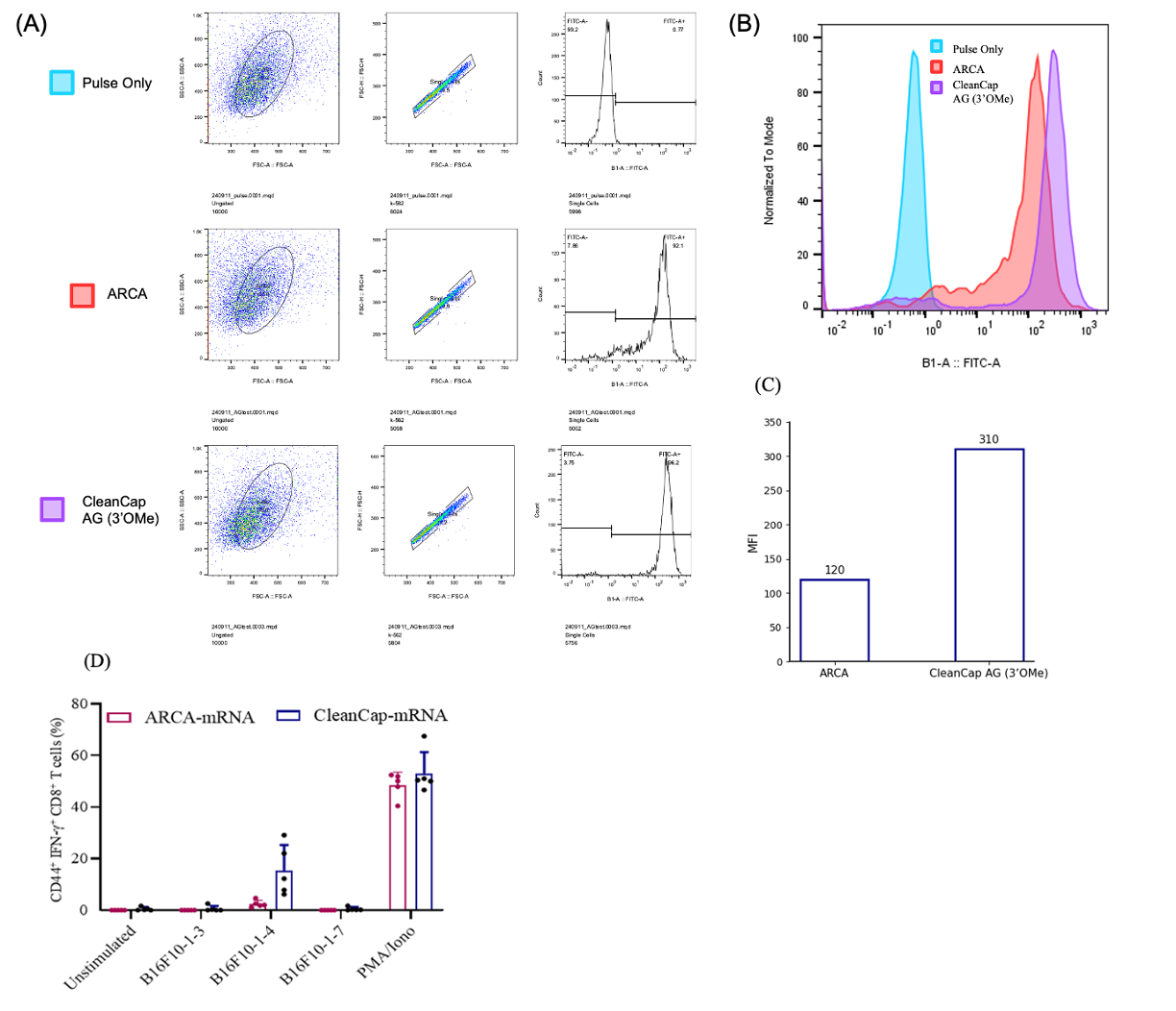
**Supplementary Figure S1 | Exploratory comparison of mRNA cap structures using EGFP reporter expression and antigen-specific CD8⁺ T-cell response assays.**

EGFP-encoding IVT mRNAs were generated using either ARCA or CleanCap® AG (3′-O-Me) cap analogs and introduced into K-562 cells by electroporation. EGFP expression was evaluated 24 h after electroporation by flow cytometry.

(A) Representative flow cytometry gating strategy for EGFP expression analysis. Live cells were selected based on FSC-A/SSC-A profiles, followed by single-cell gating using FSC-H/FSC-A. EGFP-positive cells were identified using FITC-A histogram profiles.

(B) Representative FITC-A histogram overlay showing EGFP expression in pulse-only, ARCA-capped EGFP mRNA, and CleanCap® AG (3′-O-Me)-capped EGFP mRNA conditions. Pulse-only cells were used as the negative control for EGFP gating.

(C) Quantification of EGFP expression based on mean fluorescence intensity. MFI was used as the reporter expression readout for comparison of ARCA- and CleanCap® AG (3′-O-Me)-capped EGFP mRNAs.

(D) Antigen-specific CD44⁺IFN-γ⁺ CD8⁺ T-cell responses following ex vivo peptide restimulation of splenocytes from mice vaccinated with ARCA- or CleanCap® AG (3′-O-Me)-capped mRNA. PMA/ionomycin was used as the positive control.

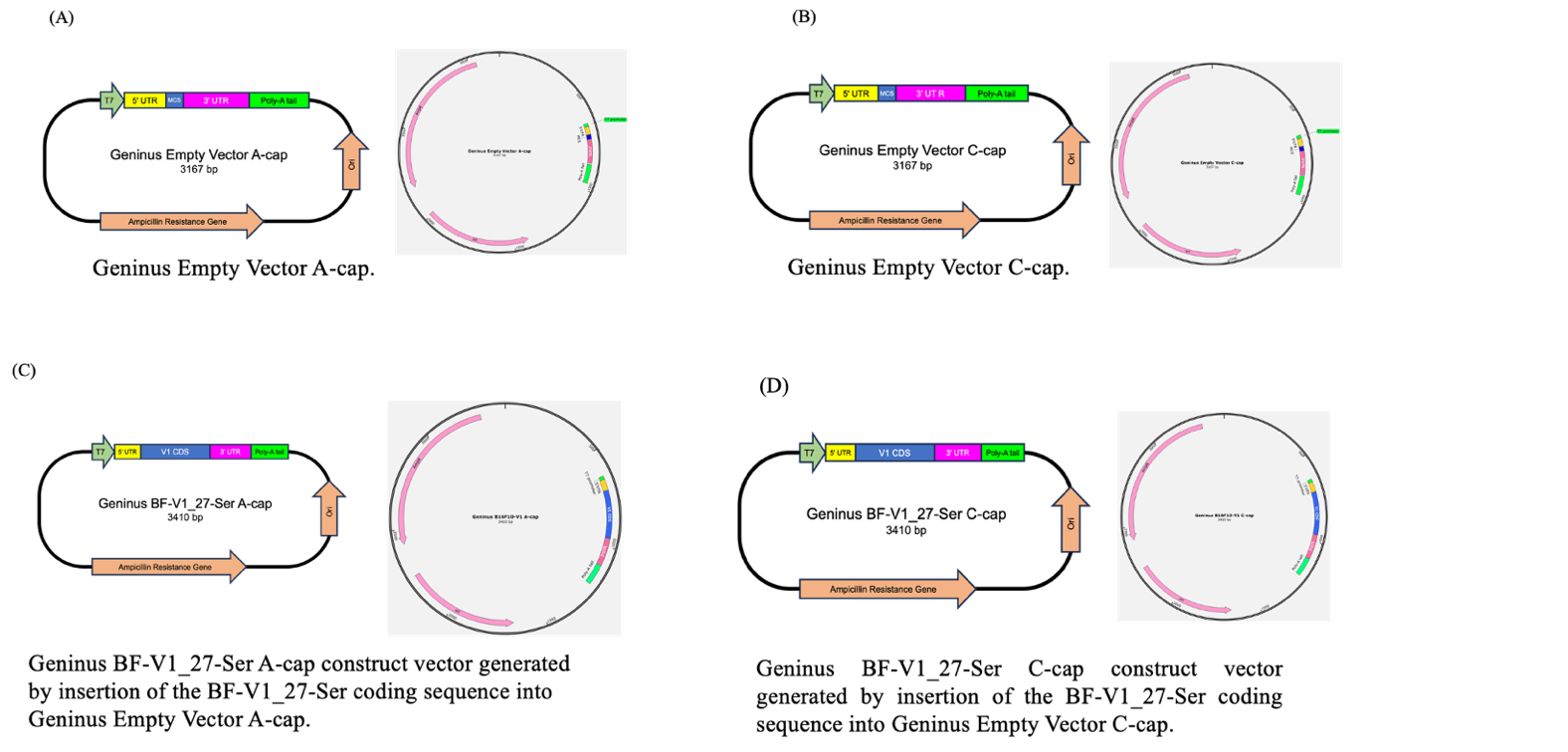

**Supplementary Figure S2 | Schematic maps of Geninus in-house vector backbones and BF-V1_27-Ser construct vectors used for IVT template preparation.**

(A, B) Schematic maps of the Geninus Empty Vector A-cap and Geninus Empty Vector C-cap backbones, respectively. Each empty vector backbone contains a T7 promoter, 5′ untranslated region (UTR), multiple cloning site (MCS), 3′ UTR, poly(A) tail, ampicillin resistance gene, and bacterial origin of replication.

(C, D) Schematic maps of the Geninus BF-V1_27-Ser A-cap and Geninus BF-V1_27-Ser C-cap construct vectors, respectively, generated by insertion of the BF-V1_27-Ser coding sequence into the corresponding Geninus empty vector backbone. The resulting construct vectors contain the T7 promoter, 5′ UTR, BF-V1_27-Ser coding sequence, 3′ UTR, poly(A) tail, ampicillin resistance gene, and bacterial origin of replication. The labels “A-cap” and “C-cap” indicate Geninus internal vector backbone variants used for IVT template preparation and do not refer to the ARCA or CleanCap® cap analogs used during in vitro transcription.
